# Deciphering the determinants of recombinant protein yield across the human secretome

**DOI:** 10.1101/2022.12.12.520152

**Authors:** Helen O. Masson, Chih-Chung Kuo, Magdalena Malm, Magnus Lundqvist, Åsa Sievertsson, Anna Berling, Hanna Tegel, Sophia Hober, Mathias Uhlén, Luigi Grassi, Diane Hatton, Johan Rockberg, Nathan E. Lewis

## Abstract

Mammalian cells are critical hosts for the production of most therapeutic proteins and many proteins for biomedical research. While cell line engineering and bioprocess optimization have yielded high protein titers of some recombinant proteins, many proteins remain difficult to express. Here, we decipher the factors influencing yields in Chinese hamster ovary (CHO) cells as they produce 2165 different proteins from the human secretome. We demonstrate that variation within our panel of proteins cannot be explained by transgene mRNA abundance. Analyzing the expression of the 2165 human proteins with machine learning, we find that protein features account for only 15% of the variability in recombinant protein yield. Meanwhile, transcriptomic signatures account for 75% of the variability across 95 representative samples. In particular, we observe divergent signatures regarding ER stress and metabolism among the panel of cultures expressing different recombinant proteins. Thus, our study unravels the factors underlying the variation on recombinant protein production in CHO and highlights transcriptomics signatures that could guide the rational design of CHO cell systems tailored to specific proteins.

## Introduction

Roughly a third of the human protein coding genome encodes secreted and membrane proteins that mediate virtually all interactions of a cell with its environment ^1^, and whose enzymatic activity regulates a diverse range of vital organismal functions. The human secretome project (HSP) ^2,3^ has comprehensively characterized this important subset of the human proteome as a resource for drug discovery and development. The fundamental roles in signaling and organismal homeostasis make these secreted proteins appealing candidates for the biopharmaceutical industry.

To recombinantly produce many biopharmaceuticals, Chinese hamster ovary (CHO) cells are the preferred mammalian expression system because of their scalability and compliance with human post-translational modifications (PTMs) ^4,5^. To systematically measure the potential of CHO cells to produce these pharmaceutical targets, an effort to express the entire human secretome recombinantly in CHO was initiated as a companion project to the HSP. Efforts were made to express 2189 secreted human proteins using the Icosagen QMCF CHO cell line (Icosagen Cell Factory OÜ), which allows for episomal extended transient protein expression. Almost 1,300 proteins have been successfully produced and purified in the cell line using the HSP standardized high throughput pipeline ^6^. We observe that the amounts of protein produced are highly variable; only 59% of the human secretome could be successfully expressed in CHO above the quality threshold. Furthermore, among the proteins that passed quality checks, titers differed by several orders of magnitude depending on the protein (Fig. 1a). This prompted us to ask the key question: what factors account for the vast variation observed in recombinant protein production in CHO? Answers to this question are of great interest in the biopharmaceutical industry and researchers across fields who study mammalian proteins, providing guidance to the rational design of recombinant protein-producing CHO cell lines.

**Figure 1.**
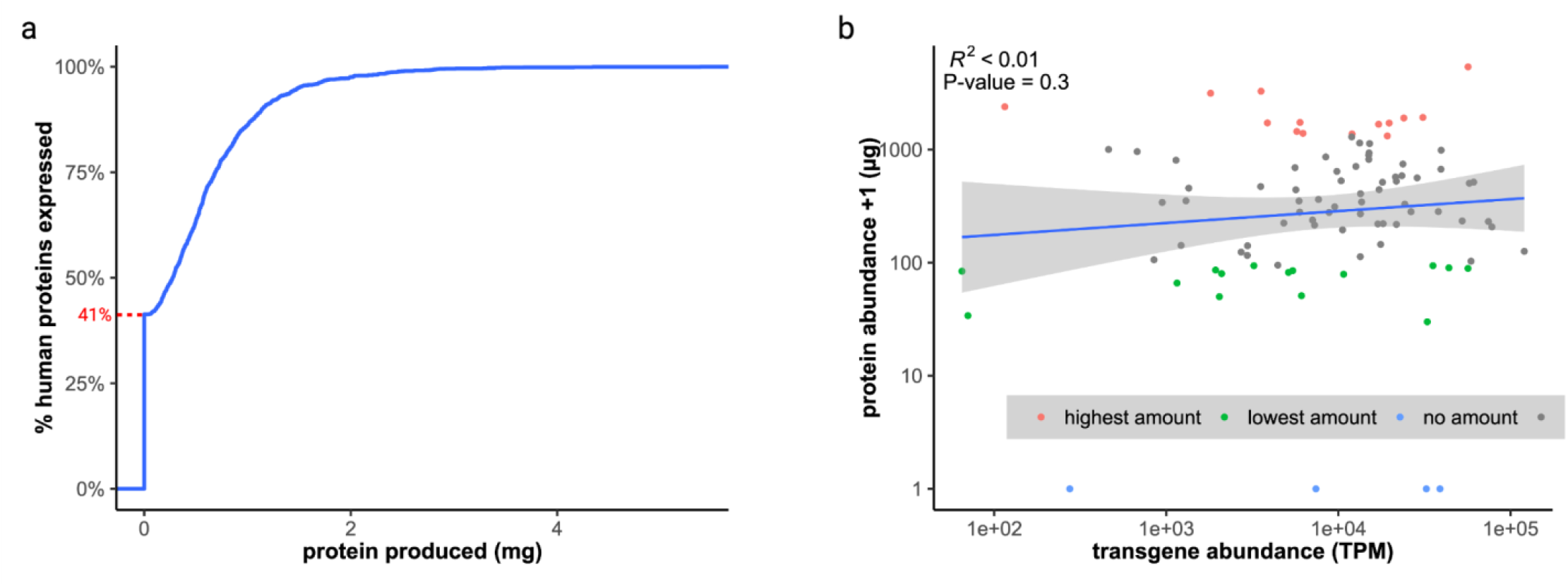
Production of the human secretome in CHO. **a)** Cumulative distribution of target protein produced for the 2165 recombinant proteins expressed using the human secretome high-throughput production pipeline. Approximately 41% (red line) of the proteins failed to produce, while the amount of recovered protein for remaining cells varied between 0.44-5.38mg. **b)** Relationship between transgene abundance (TPM) and amount of secreted protein (μg). The CHO cell line was unable to produce any recoverable product for 4 of the selected recombinant proteins (blue), while cells with the top 15 highest and lowest yields are colored in red and green respectively. Cells expressing the remaining proteins are shown in gray.

To understand the determinants of protein titers, we analyzed the expression of 2165 CHO-produced secreted human proteins (filtered set from the 2189 HSP proteins, see Methods), and conducted RNA-Seq on a representative subset of 95 CHO cell cultures, each expressing a different recombinant protein, along with the non-producing Icosagen QMCF host cell line. Here we aim to quantify the relative contribution of three major factors that influence the production and secretion of recombinant proteins. First, we modeled the relationship between transgene mRNA levels and protein yield to quantify the variability explained by transgene transcript abundance. Second, we curated hundreds of protein features and applied machine learning to identify the most important protein attributes contributing to variation in productivity. Lastly, we used transcriptomic profiles to quantify the variability explained by host cell expression signatures. We further identify specific processes associated with ER stress and metabolism that are strongly associated with the ability of cells to produce recombinant protein.

## Results

### Recombinant protein expression in CHO varies extensively

We analyzed the productivity of 2165 proteins from the HSP study and investigated the distribution of target products (Fig. 1a). Only 59% of the secretome could be successfully expressed by CHO cells above the quality threshold, determined by a combination of WB analysis, SDS-PAGE, and MS/MS at various time points^6^. Furthermore, among the proteins that passed quality checks, titers differed by several orders of magnitude depending on the protein (Fig. 1a). To enable deeper characterization of the library of CHO cells producing the human secretome, we selected a subset of 95 cell cultures each expressing a unique recombinant protein. This included high (n=15), low (n=15), and failed producers (n=4), along with 61 additional cultures wherein the produced protein varied in size and composition. We also included the wild-type (WT) Icosagen QMCF CHO-S host for comparison. This panel of 96 cell cultures were subjected to RNA-Seq, which quantified the mRNA abundance for the transgenes encoding the human secreted proteins (Supplementary Data 1), along with the endogenous CHO genes. The transgenes, as defined by their recombinant sequences, consistently take up ∼3% of the entire transcriptome, making it one of the most highly expressed genes in most samples.

### Variation in recombinant protein yield cannot be explained by transgene mRNA abundance

Some studies report that transgene mRNA levels can be limiting for secreted protein titers ^7,8^. To evaluate if the variation in protein production in our panel of cells can be explained by transgene mRNA levels, we modeled the relationship between transgene levels and protein yield using linear regression. Across the 95 RNA-sequenced recombinant protein expressing cell cultures, we found that transgene mRNA levels explained less than 1% of the variance in protein titer (Fig. 1b). This correlation pales in comparison to other studies which report numbers closer to 40% for endogenous genes in mammalian cells across various conditions ^9–12^, likely due to the high mRNA expression achieved in the QMCF system. We conclude that adequate transgene mRNA is produced in these cells, and mRNA abundance is likely not the limiting factor. These results suggest an alternative bottleneck in the production of difficult to express proteins within the HSP panel of proteins.

### A comprehensive set of 218 features describing the HSP proteins

Since transgene mRNA levels do not appear to limit recombinant protein production in our system, we wondered how protein-specific features contribute to the variability in protein yield. To test this we curated a comprehensive set of 218 protein features as potential predictors of abundance of the 2165 HSP proteins. These features were classified into three main categories: i) experimental abundance, ii) sequence features, and iii) biophysical features (Table 1). Experimental abundance features measure the expression of the protein in other systems including various human tissues, other species, and the expression of the endogenous protein in CHO. Sequence features encompass protein attributes linked to the nucleotide and amino acid sequence of the protein such as molecular weight (MW), amino acid composition (AAC), and PTMs. Lastly, biophysical features cover metrics related to protein stability, solubility, secondary structure, etc. A detailed description of all features can be found in Supplementary Data 2. The influence of these protein features on protein yield was investigated using correlation and machine learning methods.

**Table 1.** Protein features and their sources.

| Feature Type | Feature Group | # Features | Description | Source/Software Packages |
| --- | --- | --- | --- | --- |
| Experimental abundance | Production in mouse | 18 | Protein and mRNA copy numbers, half-lives, transcription rates and translation rate constants in mouse fibroblasts | 10.1038/nature10098 |
|  | Production in yeast | 1 | Production yield of fusion proteins with fractions of human secretome in yeast. | 10.1093/bioinformatics/btx207 |
|  | Human tissue expression | 17 | Secretome expression in various human tissues | GTE<sub>x</sub> |
|  | Human tissue protein level | 19 | Protein level across different human tissues | HPA |
|  | Endogenous expression of CHO ortholog | 6 | Endogenous expression of CHO ortholog under various conditions | this study |
|  |  |  |  | 10.1038/srep40388 |
| Sequence features | Molecular weight | 1 | Molecular weight of protein | this study |
|  | Post-translational modifications | 29 | Number of post-translational modifications normalized with respect to sequence length | 10.1371/journal.pone.0063284<br>iPTMnet<br>ScanPROsite |
|  | AA composition | 20 | Amino acid composition (AAC) | this study |
|  | AA composition correlation with CHO | 22 | Correlation of AAC with AAC in native CHO cells |  |
|  | AA class composition | 30 | Global percentage of various AA classes | Peptides<br>protr |
|  | AA class transition | 21 | Percent frequency of transitions between pairs of AA classes | protr |
|  | RNA secondary structure | 3 | RNA minimum free energy (MFE), normalized ensemble free energy (EFE), and MFE normalized with respect to sequence length | RNAfold |
| Biophysical features | Stability | 4 | Stability, instability, and aliphatic indices | Peptides<br>ProtParam<br>ProTstab |
|  | Solubility | 7 | Isoelectric point, net charge, percent solubility, and grand average of hydrophobicity (GRAVY) | Peptides<br>ProtParam<br>Protein-Sol |
|  | PPI potential | 1 | Potential protein protein interaction index. | Peptides |
|  | Secondary structure | 11 | 3- and 8-category predictions of protein secondary structure | Scratch |
|  | Relative solvent accessibility | 8 | Solvent-accessible fraction, percent hydrophobic and hydrophilic solvent-accessible residues, mean accessibility score, and GRAVY of inner and outer residues | Scratch |

### MW, AAC, and N-linked glycosylation have the greatest effect on protein titers

The importance of individual protein features was quantified using Spearman correlation (Table 2). Using the subset of proteins that passed quality control and produced at detectable levels, we found that MW had the strongest correlation (R=0.26) with protein yield (μg). This unexpectedly suggests that higher molecular weight proteins were easier to produce. To understand this further, we binned the proteins by MW and observed that the significant correlation only holds true for low MW proteins (Supplementary Fig. 1-2). A significant drop in correlation was observed once the protein surpassed 2500-3500 Da, suggesting a sort of size threshold below which protein size becomes difficult to produce efficiently. We also observed a significant correlation between AAC of cysteine and protein yield (R=-0.23). Cysteines are involved in the formation of molecular architecture-mediating disulfide bonds, which also showed a similar relationship with protein yield (R=-0.14). This negative relationship suggests that recombinant proteins containing a high proportion of cysteines and disulfide bridges tend to produce less efficiently.

**Table 2.** Correlation between protein features and protein yield (μg)

| Feature Group | Feature | Correlation with protein yield (µg) |
| --- | --- | --- |
| Molecular weight | MW (Da) | 0.256*** |
| AA composition | AA. comp C | -0.225*** |
| AA composition correlation with CHO | AA. comp correlation with native CHO | 0.198*** |
|  | AA. comp correlation with essential CHO | 0.166*** |
| AA class composition | AA. comp med volume | 0.139*** |
| Post-translational modifications | N-linked glycosylation | 0.159*** |
|  | Disulfide bonds | -0.140*** |
| Secondary structure | Coil | -0.189** |
| Relative solvent accessibility | Mean accessibility score | -0.199*** |
|  | Percent hydrophobic solvent-inaccessible residues | 0.188** |
|  | Percent hydrophobic solvent-accessible residues | -0.144** |
| Stability & Solubility | Net charge | -0.169*** |
|  | Grand average of hydropathicity | 0.166*** |
|  | Isoelectric point | -0.138*** |
|  | Instability index | -0.130*** |

List of selected protein features amongst predictors with the strongest Spearman correlation coefficient with protein yield. Significance values were adjusted using false discovery rate (FDR) method to correct for multiple testing: \*P ≤ 0.01, \*\*P ≤ 0.001, \*\*\*P ≤ 0.0001.

To further understand the complex relationship between protein features and yield, we generated descriptive regression and classification models of recombinant protein production in CHO using machine learning (ML). Regression algorithms using the subset of quantifiable proteins that passed quality control provides insight into features hindering lowly expressed proteins. On the other hand, classification models using the pass/fail status of proteins can elucidate features preventing the production of proteins. Protein features were filtered and preprocessed before serving as predictors in both regression and classification pipelines, each of which produced 8 unique models (see Materials and Methods). Predictor variable (i.e. protein feature) importance for each model was ranked, and the consensus among the top 10 predictors for each model was evaluated (Fig. 2a-b). All 8 regression models ranked MW and AAC of cysteine amongst the top 10 most important features affecting protein yield. This supports the correlation analysis which identified these same two features as having the strongest correlation with protein abundance. Furthermore, the best performing regression model ranked these predictors as the most important features affecting protein yield (Fig. 2c). Our classification models using the pass/fail status of proteins showed increased consensus among important protein features. Among the universally consented features were N-linked glycans, which are critical for folding and quality control of glycoproteins, specifically through the calnexin/calreticulin cycle ^13,14^. When we set the failed samples to zero titer and performed a correlation analysis, we found a significant positive correlation between N-linked glycosylation and yield (R=0.26) (Supplementary Table 1), indicating that proteins with increased N-linked glycosylation tend to express better.

**Figure 2.**
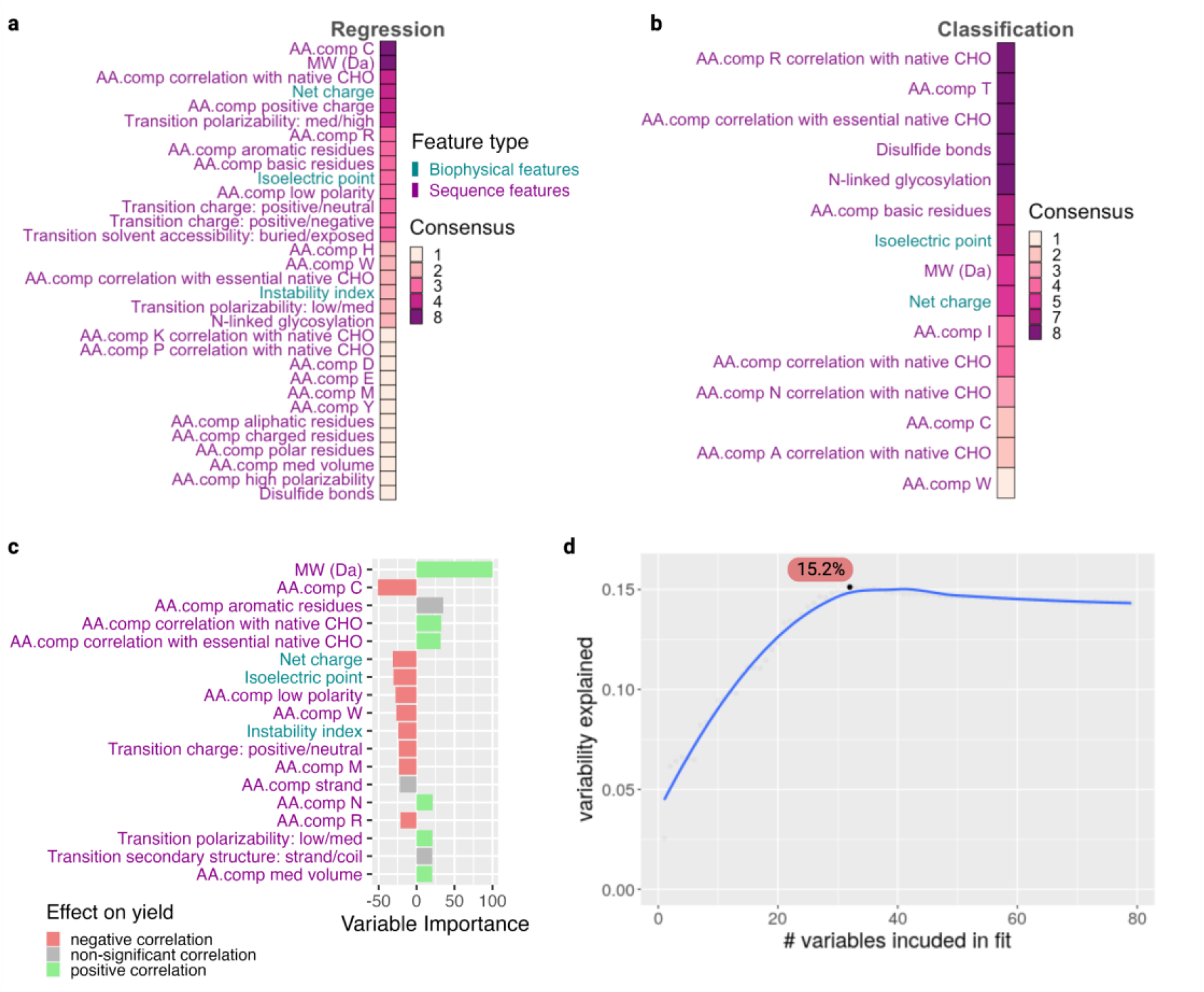
Protein-specific features affect recombinant protein yield. **a-b)** Compilation of the top 10 most important features identified in the 8 regression (a) and 8 classification (b) models. A consensus of 8 indicates that the feature was identified as an important feature in all 8 models. Regression models showed lower consensus highlighting a total of 32 features, only 2 of which showed up in the top 10 features of all 8 models (consensus=8). However, the classification models showed higher consensus highlighting a total of 15 features, wherein a third (5) of them have been deemed highly important in all 8 models (consensus=8). **c)** Bar graph showing the most influential protein features identified in our best performing regression model. Variable importance measures have been scaled to have a maximum value of 100, and their directional effect on yield has been inferred and colored based on the feature correlation with protein titer. **d)** Variability in protein titers explained by protein features was determined by sequentially adding protein features to a linear regression model and calculating the percent variability explained by the set of features. AA comp: amino acid composition; MW: molecular weight. A detailed description of each protein feature can be found in Supplementary Data 2.

### Protein features account for ∼15% of the variability in recombinant protein yield

Protein features, in particular sequence features, clearly affect CHO’s ability to successfully produce recombinant protein and may help inform recombinant protein candidate selection or design for future production runs. To quantify the variability in protein yield that can be explained by protein features, we sequentially added the ranked features of the best performing regression model to a linear model fit and calculated the fraction of variance explained by the model (Fig. 2d). The explained variance peaks at approximately 15% when 32 protein features are included. While significantly greater than the variability explained by transgene mRNA abundance, protein features only account for a fraction of the variability in protein titers. Together these results suggest that protein features are not the most important factor limiting recombinant protein production in CHO.

### Transcriptomic signatures can account for the majority of variation in protein titers

Targeting protein features to enhance titers is typically undesirable as the features can be integral to protein function. We therefore investigated how transcriptomic determinants in the host cell impact protein yield. Principal component analysis of the 96 RNA-Seq samples (Supplementary Data 3) clearly shows that the non-producing cells, including WT, are transcriptional outliers compared to the cells producing recombinant protein (Fig. 3a). The first principal component (PC1) accounts for approximately 19% of transcriptome variability, and separates successfully producing cells from those that failed to produce any recombinant protein. LOC100754005, one of the top 5 influential genes with a negative loading on PC1, encodes an ortholog of the PRPF8 gene (Pre-mRNA-Processing-Splicing Factor 8) which serves as a component of the spliceosome critical for pre-mRNA processing. We find that higher expression of this gene differentiates the productive cell lines from the non-producing outliers. Interestingly, previous work comparing the proteome of various CHO host cells revealed an up-regulation of PRPF8 in the high producing cell lines and alluded to its contribution to the high production of biopharmaceuticals in CHO ^15^.

**Figure 3.**
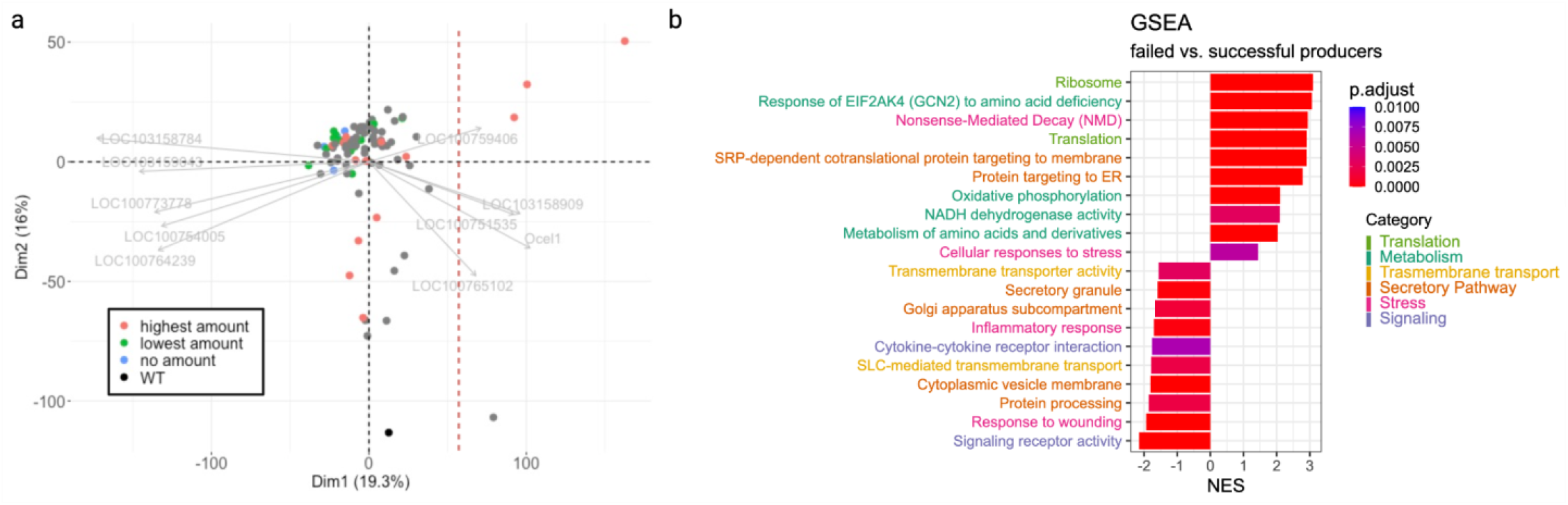
Non-producing cell lines are transcriptional outliers. **a)** Principal component analysis (PCA) of transcriptomics data. Top 5 positive and negative contributing genes to the first principal component (PC1) shown in light gray. Dashed red line shows a clear division between the cells capable of producing recombinant proteins (red, green, and gray) and cells that failed to produce any detectable protein (blue). **b)** Results from a gene set enrichment analysis (GSEA) performed between the failed producers and the cells that successfully produced protein. Terms with a positive normalized enrichment score (NES) are enriched by genes overexpressed in the non-producers, while terms with a negative NES are enriched by genes overexpressed in the producers.

To gain additional insights into biological pathways and processes characteristic of the non-producers, we conducted Gene Set Enrichment Analysis (GSEA) ^16,17^ between the failed producers and the cells that successfully produced protein (Fig. 3b). Unsurprisingly, we saw signs of cell stress (pink terms) in both groups, likely due to the burden of overexpressing foreign protein. Additionally, we found that the failed producers upregulated genes involved in translation (green terms) and oxidative phosphorylation, and showed signs of amino acid deficiency (teal terms). We also observed increased activity in the early stages of protein secretion (i.e targeting to the ER) in the failed producers, and depletion in later portions of the secretory pathway (i.e. Golgi subcompartments, vesicle membranes, and secretory granules) compared to the producers (orange terms). Furthermore, the successful producers show increased transmembrane transport (yellow terms), potentially alleviating the burden of amino acid deficiency.

To quantify the variability in protein yield explained by transcriptomic cell signatures, we conducted multiple linear regression on the principal component loadings. Using the first three principal components, which account for 44% total variation of the transcriptome, we found that host cell gene expression signatures could account for 75% of the variability seen in protein yield. Even though our panel of cells come from a single clonal cell line, the expression of different transgenes is clearly impacting the cells in a protein-specific manner.

### Cells respond differently to ER Stress

Our GSEA analysis alluded to significant differences in secretory pathway activity. To better understand the protein-specific secretory pathway signatures within our panel of cells, we calculated activity scores (see Materials and Methods) for 13 secretory pathway functions (Supplementary Data 6). Activity scores for the 95 recombinant protein expressing CHO cells were normalized to express the change in pathway activity with respect to the WT host cell (Fig. 5a).

ER calcium homeostasis was the most highly increased function across recombinant protein expressing cells regardless of productivity, suggesting that overexpression of heterologous proteins in CHO triggers a general imbalance in ER calcium homeostasis. Maintaining proper Ca^2+^ levels within the ER is vital for virtually all ER-supported functions, and disruption of these levels activates ER stress and UPR^18^. In fact, an in-depth analysis of cellular response to stress (Supplementary Results) showed activation of many ER stress response genes among the panel of cells. In particular, results show a depletion in all three branches of UPR signaling and signs of increased ubiquitin-mediated proteasomal degradation (ER-associated degradation; ERAD) in the failed producers.

Protein folding in particular is a common bottleneck in recombinant protein production, and the accumulation of improperly folded proteins can also trigger ER stress. However, the upregulation of protein folding genes is associated with greater protein production ^19–22^. In line with these findings, we observe a mild yet significant positive correlation between protein folding activity and protein yield (r=0.21, pval=0.05, Supplementary Data 12). Furthermore, the stress analysis (Supplementary Results) identified several genes involved in disulfide bond formation and protein folding including HYOU1 (hypoxia up-regulated 1), ERO1A (endoplasmic reticulum oxidoreductase 1 alpha), and PDIA3 (protein disulfide isomerase family A member 3) upregulated alongside the stress response in the successfully producing cells. Altogether, these results suggest that the productive cells respond to ER stress better than the failed producers.

### N-linked glycosylation and ERAD are strong determinants of protein yield

Clustering the 95 recombinant protein expressing cells based on secretory pathway activity on transcriptional level revealed 4 distinct groups (Fig. 4a). One cluster consists of a single failed producer (CCL20) that shows dramatic decreases in activity across all secretory pathway functions, while the other 3 clusters show unique secretory pathway footprints. Cluster 1 shows little to no change in the majority of secretory functions, cluster 2 is characterized by a general increase in activity across subsystems, and cluster 3 is characterized by a general decrease in subsystem activity. Similar to the single non-producing outlier which showed a dramatic decrease in secretory pathway activity, the remaining 3 non-producing cell lines also show decreased secretory pathway activity and belong to cluster 3. The cells in each cluster show a range of productivity, suggesting these secretory pathway footprints do not define a cell’s ability to successfully produce and secrete recombinant protein. Of particular interest was cluster 3, which showed low activity across all secretory functions. Given that some of the highest producers fall within this cluster, overall high secretory pathway activity is not required for high protein yield. However, when calculating pathway activity scores we lose gene-specific granularity. Therefore we wondered if there are sets of genes that drive the high protein production seen in certain cells of cluster 3.

**Figure 4.**
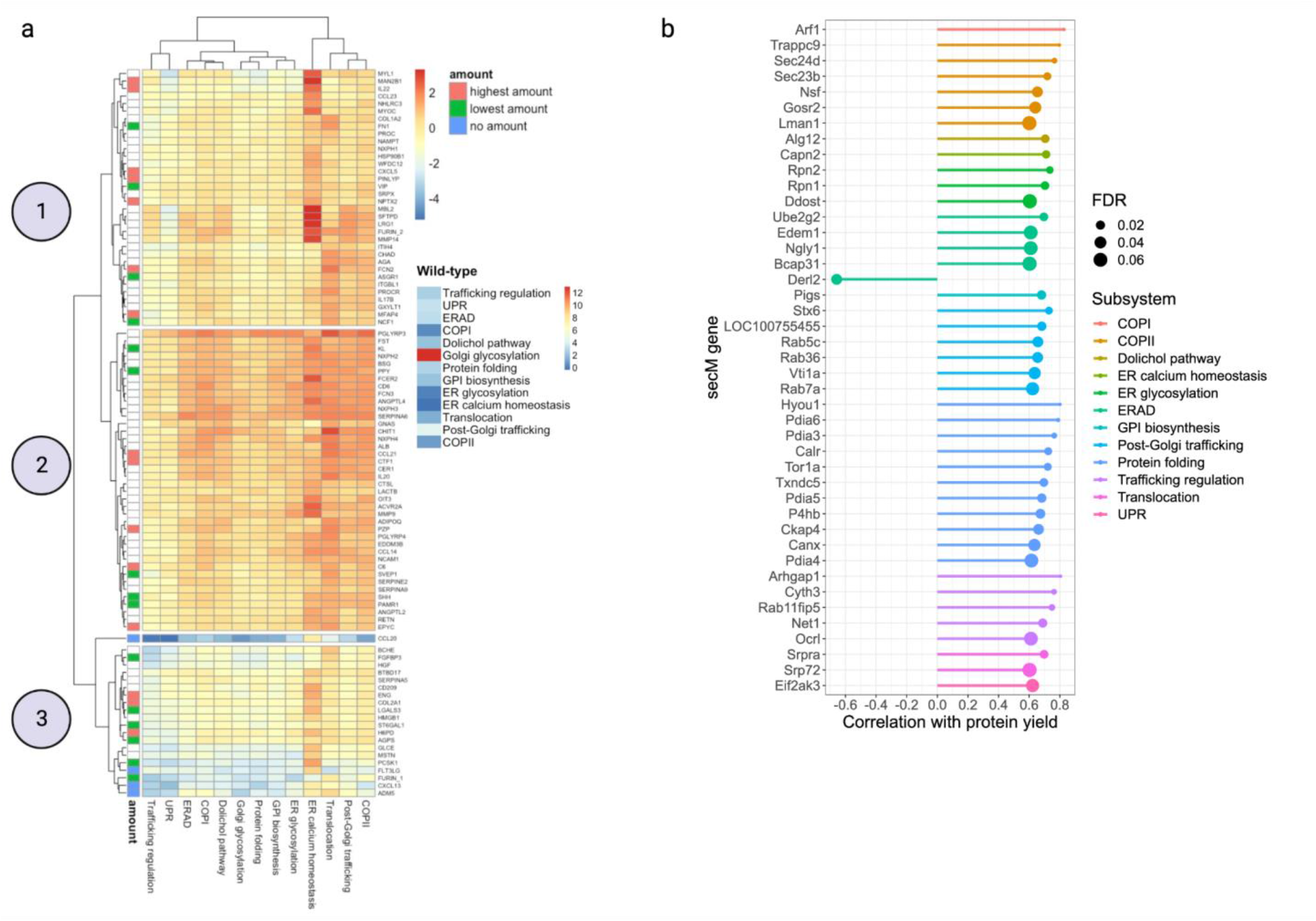
Secretory pathway cell signatures. **a)** Clustered heatmap of the normalized change in secretory pathway activity compared to WT for each of the 95 recombinant protein expressing cells. Highlighted here are 3 clusters that show distinct secretory pathway footprints. Cells are annotated according to the amount of protein they produce: no protein (blue), highest yield (red), and lowest yield (green). Raw scores for WT are shown to the right. **b)** Lollipop plot showing the significant correlations between secretory pathway genes and protein abundance amongst the cells in cluster 3.

To understand which genes drive high production of recombinant protein in cluster 3, we calculated correlations between individual secretory pathway genes and protein abundance for the cells of the cluster (Supplementary Data 7). We identified 43 secretory machinery genes that showed significant correlation (|r|>=0.6; false discovery rate (FDR)<= 0.1) with protein abundance (Fig. 4b). One set of positively correlated genes was particularly interesting: Alg12 (Alpha 1,6 Mannosyltransferse), Rpn1 (Ribophorin 1), Rpn2 (Ribophorin 2), and Ddost (dolichyl-diphosphooligosaccharide-protein). While these genes belong to different subsystems, dolichol pathway and ER glycosylation, they are involved in the same integral process of N-linked glycosylation. Alg12 encodes a glycotransferase involved in the assembly of the dolichol-PP-oligosaccharide precursor required for N-linked glycosylation. Rpn and Ddost encode proteins of the oligosaccharide transferase complex (OST complex), which catalyzes the first step of N-linked glycosylation – the transfer of the pre-assembled N-glycan from the dolichol lipid carrier to the client protein. Given the importance of N-linked glycosylation in protein folding and quality control within the ER, it is reasonable to believe that genes involved in this step are critical for efficient protein secretion.

Only a single gene, Derl2 (Derlin 2), was negatively correlated with protein yield. The derlin genes encode components of ERAD machinery, where they participate in the retro-translocation of unfolded and misfolded proteins from the ER to the cytosol for proteasomal degradation ^23,24^. Interestingly, derlins also function in ER-stress induced pre-emptive quality control (ERpQC)^25,26^. During ER stress, Derlin is recruited to the translocon and signal recognition particle receptors and participates in the selective attenuation of translocation of newly synthesized proteins into the ER, rerouting them to the cytosol for proteasomal degradation. The downregulation of this ERpQC mechanism allows proteins to enter the ER and interact with protein folding chaperones, increasing the chances of protein production and secretion. We used linear regression to quantify how much of cluster 3’s variability in protein abundance could be attributed to the 5 aforementioned genes: Alg12, Rpn1, Rpn2, Ddost, and Derl2. Due to overlapping biological functions, the expression of Rpn1, Rpn2, Ddost, and Alg12 are highly correlated, therefore to avoid multicollinearity we only included the expression of Derl2 and Alg12. The resulting model could explain an astonishing 87% of cluster 3’s variability in protein yield. These results suggest that Alg12 and Derl2 may be good engineering targets, especially for cell lines with overall low secretory pathway activity.

### Failed producers are metabolically less active

Recombinant protein production is energy intensive with increased raw material demands, thus inducing significant alterations in host cell metabolism. Consequently, many cell line engineering efforts have targeted metabolism to enhance recombinant protein production ^27^. To identify metabolic variation within our panel of cells, we implemented the CellFie tool ^28^, which quantifies metabolic task activity from omics data (Supplementary Data 8). We identified 79 core metabolic tasks active in all cells, 27 tasks inactive across all cells, and 79 tasks with differential activation (Fig. 5a). Many differentially active tasks are involved in amino acid and carbohydrate metabolism. When looking at the 79 tasks showing differential activation across our panel of CHO cells, the non-producers showed on average 33% active metabolic tasks, while the highest and lowest producers showed 66% and 58%, respectively (Fig. 5b), suggesting the non-producers are metabolically less active compared to the producing cells.

**Figure 5.**
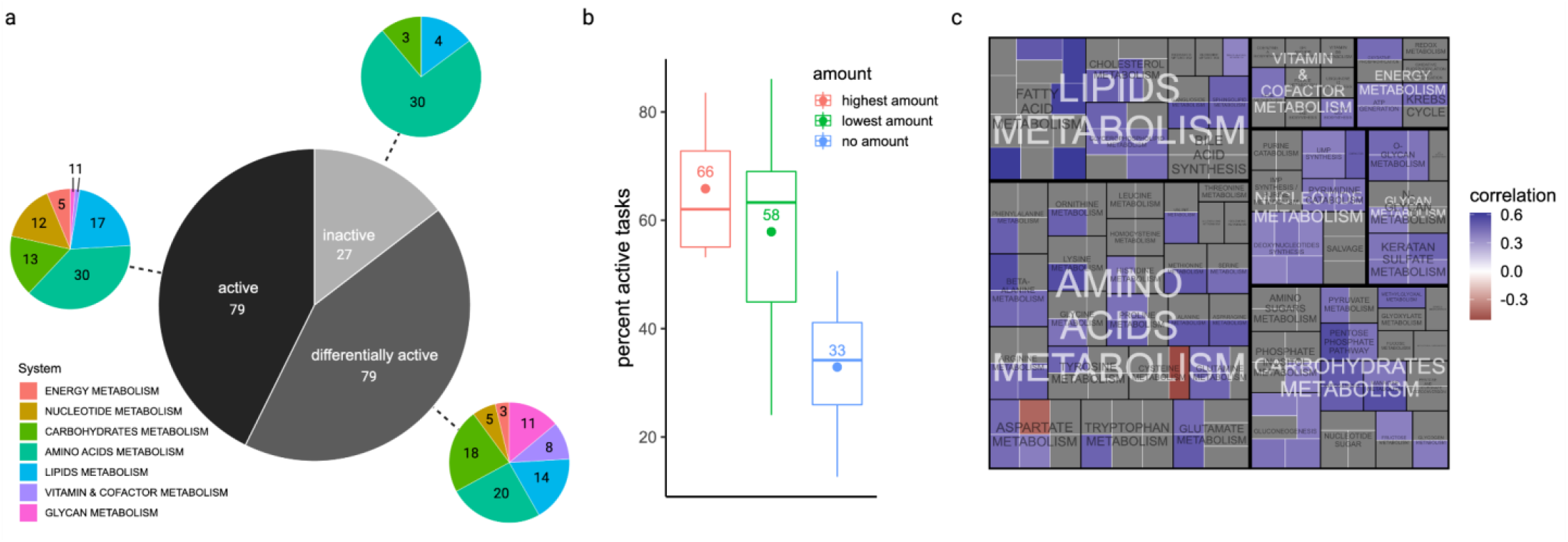
Metabolic cell signatures. **a)** Proportion of tasks that are active, inactive, and differentially active among the 95 recombinant protein expressing cells. Also displayed are the proportions of tasks falling within each subsystem. **b)** Boxplot showing the percentage of active metabolic tasks among the different productivity groups. **c)** Treemap of CellFie metabolic tasks organized into systems and subsystems. Each square represents a single metabolic task which is colored according to significant correlation with protein yield among the high and low producing cell lines.

### Increased fatty acid metabolism in the high producers

To further understand the metabolic differences, we used the quantitative form of metabolic scores to characterize the relationship between individual tasks and protein yield. Several metabolic tasks showed significant correlations (FDR<=0.1) with protein abundance among the subset of high and low producers (Fig. 5b; Supplementary Data 9). The majority of tasks show positive correlation with protein abundance, further suggesting that higher metabolic activity facilitates recombinant protein production. We found that the metabolic tasks with the largest and most significant correlation with protein yield among the subset of high and low producers are involved in fatty acid (FA) metabolism. In particular, we observe a strong positive correlation with synthesis of several FAs: palmitoleate synthesis (R=0.62), palmitate synthesis (R=0.61), synthesis of palmitoyl-CoA (R=0.59), arachidonate synthesis(R=0.59), and synthesis of malonyl-coa (R=0.51). FAs have a diverse range of important cellular functions including critical structural components of cell membranes. Cells modulate the FA composition of the cell membrane under challenging conditions to regulate membrane fluidity^29^. Increased activity in FA metabolism may be a signature characteristic of high recombinant protein production, given its importance in the size and function of the endomembrane system and the secretory pathway in general. Additionally, FAs can store and supply energy to cells. Our results revealed a positive correlation between the stress response energy-producing FA oxidation gene ACAA2 (acetyl-CoA acyltransferase 2) and protein yield (Supplementary Results, Supplementary Fig. 3b). In combination with the observed overall increase in FA metabolism, these results could suggest that the high producing cells are using FA metabolism to provide a beneficial pool of energy to meet the demands of high recombinant protein production.

### Cysteine depletion and oxidative stress in the poor producers

We found only two tasks, conversion of aspartate to beta-alanine and synthesis of taurine from cysteine, showed a negative relationship with protein yield (Fig. 5c). Our protein features analysis showed that our host system has difficulty producing proteins with high cysteine composition. The depletion of available cysteine from the synthesis of taurine could be further burdening the production of proteins. Furthermore, not only does this task deplete the availability of free cysteine, but there is evidence that taurine acts as an antioxidant defense by counteracting lipid peroxidation ^30,31^ which could be an indicator of increased oxidative damage.

The prevalence of oxidative stress within our panel of cells was further confirmed by our in depth analysis of cellular response to stress (Supplementary Results). Firstly, we noticed that the successfully producing cells show a more profound response to oxidative stress, upregulating almost twice as many oxidative stress response genes compared to the non-producing cells. Second, we observed that three of the genes depleted in the failed producers encode proteins belonging to the solute carrier (SLC) superfamily, supporting the negative enrichment in SLC transmembrane transport observed in the preliminary GSEA analysis. SLC7A11 (solute carrier family 7 member 11) shows the greatest depletion among oxidative stress genes in the failed cells (LFC=-1.85, FDR=5.19E-07) and is involved in the specific transport of cysteine and glutamate. The ability to mount an adequate response against oxidative stress, including enhancing the transport of cysteine, may facilitate recombinant protein production.

## Discussion

The continual discovery of new biologics is accompanied by pressure to establish novel methods and technologies for enhancing quality and productivity. CHO cells dominate biotherapeutic protein production and are extensively used in mammalian cell line engineering research due to their human-compatible PTMs and adaptability to suspension-growth culture in chemically-defined media. However, many proteins struggle to express well or at all in this non-native environment. The Human Secretome Project demonstrated that even standard human proteins can be difficult to produce. This large data set of heterologous protein expression in the most popular biopharmaceutical expression host represents an attractive resource that can be leveraged to understand why CHO cells produce some proteins better than others. In particular, this study was designed to illuminate and quantify the factors contributing to this variation in productivity to help guide the rational design of protein-specific CHO cell systems. Here we found that transgene mRNA levels were expressed at consistently high levels and cannot explain the variability in protein yield (<1%; Fig. 1b), allowing us to identify other factors as the main drivers in protein yield.

Using statistical and ML methods, we systematically quantified how 218 protein features affect the efficacy of protein production in CHO. Both correlation and ML analyses implicate MW and cysteine AAC as important protein features influencing efficient production in CHO (Table 2; Fig. 2). We observed a MW threshold ∼2500-3500 Da below which proteins become difficult to produce efficiently (Supplementary Fig. 2). Studies have shown that protein size is the primary factor in determining folding rates and protein stability^32^. Furthermore, small proteins are more sensitive to changes in stability than larger proteins^33^. Perhaps the small proteins lack the molecular material to form sufficient stabilizing bonds resulting in poor yield. Alternatively, this observation could be due to protein detection methods where low MW proteins are vulnerable to poor retention and resolution. We also observed a negative relationship between cysteine composition and protein yield.

Cysteine residues are important to the conformational stability of a protein through the formation of disulfide bridges which occur upon oxidation of the thiol groups between two spatially proximal cysteines. However the same property that allows this stabilizing bond formation to occur also imparts intrinsic vulnerability to oxidative stress. The highly reactive nucleophilic thiol group can be reversibly or irreversibly modified and lead to dysfunctional protein ^34^. Given we found strong transcriptional signatures of oxidative stress among the panel of cells (Supplementary Fig. 3a), high cysteine composition could be introducing destabilizing non-native disulfide bonds. In fact, studies attempting to stabilize proteins by introducing artificial disulfide bridges have found that it can lead to overall protein destabilization^35–39^. Another possible explanation is that the cysteines are forming intermolecular bonds leading to protein aggregation, since aberrant protein aggregation can occur from oxidation-induced intermolecular disulfide bond formation ^40,41^. Alternatively, the production of proteins with high cysteine composition could be depleting cysteine from the system. Indeed, cysteine depletion can induce oxidative stress, ER stress, reduced viability, and lower titers in CHO bioproduction^42,43^. Lastly, we observed N-linked glycosylation as an important protein feature enhancing recombinant protein production (Table 2; Fig. 2). Heterologous protein production can be enhanced with added N-linked glycosylation sites ^44–46^ by stabilizing the protein and enhancing quality control checkpoints. While protein features seem like a promising feature that could improve protein production, overall we found the protein features tested only account for a fraction of the observed variability in protein yield (∼15%).

Ultimately, the majority of variability (75%) in protein production was explained by cell signatures in the host transcriptome. Further transcriptomic analyses of cell stress, protein secretion, and metabolism suggest that recombinant proteins impose unique burdens on the cell. It is unsurprising that overexpression of foreign proteins induces cell stress, and in particular ER Stress. Many studies have implicated the secretory pathway, specifically the ER, as a major bottleneck in recombinant protein production ^47–49^. Our results suggest that the cells that can successfully produce recombinant proteins may also better mitigate ER stress by triggering UPR signaling and increasing protein folding machinery; meanwhile, failed producers upregulate protein clearance strategies, e.g., ERAD and ERpQC. We also observed a decrease in metabolic activity in poor producers (Fig. 5), suggesting these cells cannot keep up with the increased energy and raw material demands of recombinant protein production and secretion. Other studies have reported similar metabolic restructuring when comparing cells producing secreted vs. intracellular proteins, implicating increased energy demand of the secretory pathway during recombinant protein production^50^. The strongest metabolic differences we observed involve the metabolism of FAs, which serve as integral constituents of the secretory pathway endomembrane system and as a cell energy source. Thus, lipid metabolism might enhance recombinant protein production by allowing cells to maintain lipid homeostasis in a state of dynamic lipid turnover, or provide a beneficial pool of energy to meet the demands of high recombinant protein production. Lastly, results implicate the metabolic depletion of cysteine as negatively affecting the efficient production of protein in CHO. This corresponds nicely with our observation of high cysteine composition in the poor producers. Cysteine deprivation can trigger amino acid deprivation pathways^51^ and induce mitochondrial dysfunction leading to reduced oxidative phosphorylation^43^, both of which we observed here. Furthermore the production of the antioxidant molecule taurine from cysteine could be a result of increased oxidative stress in the poor producers.

In conclusion, results here have important implications for mammalian bioproduction. The factors underlying the variability in protein production in the most popular expression host identified here can be leveraged to improve recombinant protein production in CHO^52^ and have considerable impact on the vast biologics industry. Furthermore, this study has important implications across a range of other fields as it identifies essential processes regulating protein secretion, thus impacting cell-cell interactions associated with normal and pathological processes in the human body such as development, immunology, and tissue function.

## Methods

### Human secretome production data

Protein titers for the human secretome transiently expressed in the Icosagen QMCF cell line were taken from Tegel et al^6^. We removed samples whose status is “Ongoing”, as well as samples that passed QC (Status = “Pass”) yet were missing titer information. This left us with data for 2165 different proteins of the human secretome expressed in CHO. This cleaned up version of the data can be found in Supplementary Data 10. We note that as previously reported^6^, the titers were estimated upon purification, which could influence the results if different proteins purified differently. However, all purifications relied upon the same peptide tag, thus minimizing potential biases. Here we measured single replicates for each protein. Future studies incorporating alternative purification methods and increased replicates will further strengthen analyses into the factors affecting recombinant protein secretion in CHO.

### Sequence processing and RNA-Seq quantification

Sequence data for RNA-Seq were quality controlled using FastQC and summarized with multiQC ^53^. Trimmomatic ^54^ was used to trim low-quality bases and sequencing adapters from the reads with the following parameters: LIDINGWINDOW:5:10 LEADING:15 TRAILING:10 MINLEN:36 TOPHRED33. The CHO-K1 reference genome ^55^ was extended to incorporate the transgene sequences so that the transcripts of the heterologous secretome can be quantified. Reads were then quasi-mapped to the extended CHO-K1 genome and quantified with Salmon ^56^ with default parameters.

### Quantifying effect of mRNA abundance on protein yield

Transgene mRNA abundance was plotted against total protein yield (μg) on a log scale using ggplot2 ^57^ in R ^58^. A pseudo count of 1 was added to protein abundance to account for samples which failed to produce any detectable recombinant protein. Note the sample producing IL22 was removed due to issues quantifying the transgene mRNA abundance. A linear model was fit to the data, and model estimates displayed using the ggpmisc package ^59^.

### Protein features importance

To fully characterize the properties of the human secretome dataset, we built upon the features from our pilot study ^60^ which reviewed the expression determinants of the human protein fragments used in the creation of the antibodies for the HPA project. The final compendium of curated features included 218 metrics generated from numerous resources (Supplementary Data 2). Individual predictor importance was evaluated using non-parametric Spearman rank correlation. Significance values were adjusted using FDR to correct for multiple testing. The machine learning pipelines were built using the caret package ^61^ in R. Note that the transgene mRNA level was excluded from this analysis to isolate the effect of the recombinant protein features. All features were pre-processed (normalization, removal of highly correlated variables and incomplete features). The regression pipeline generated 8 regression models: i) glmnet, ii) partial least squares, iii) averaged neural network, iv) support vector machines with radial basis function kernel, v) stochastic gradient boosting, vi) boosted generalized linear model, vii) random forest, and viii) cubist. Similarly, our classification pipeline implemented the same first 7 algorithms (i-vii), however the cubist algorithm is unique to regression, so a naive Bayes model was used for the 8th and final classification model.

As these were generated as descriptive and not predictive models of protein features, the models tended to overfit the data. To avoid reporting an inflated metric of explained variance, we used a standard linear regression fit to calculate the variability explained by protein features. We took the rank-ordered features of the best performing regression model, and sequentially added the features to the linear model fit.

### Transcriptomic determinants of protein secretion

Low count genes were filtered based on GTEx’s scheme: expression thresholds of >0.1 TPM in at least 20% of samples and ≥6 reads in at least 20% of samples. Expression values were then log transformed to reduce heteroscedasticity concerns in downstream analyses. To facilitate functional annotation, an ortholog conversion table ^62^ was used to convert CHO genes to their human ortholog. Principal component analysis was conducted using the stats package included in R and visualized in a biplot using the factoextra package ^63^. GSEA was conducted using the clusterProfiler package ^64^ to determine the significantly up- and down-regulated cellular processes associated with the first principal component. Annotations for the enrichment were obtained from GO, Reactome, and KEGG databases. A normalized enrichment score (NES) representing the GSEA statistic (Subramanian et al., 2005) was calculated to quantify the overall direction of regulation for each gene set along with an accompanying permutation p-value which has been adjusted to correct for multiple testing.

We used multiple linear regression to quantify the overall variability in protein yield explained by transcriptomic cell signatures. To avoid the curse of dimensionality (more genes in the transcriptome than samples in our data set), we used loadings from the first 3 principal components, which account for 44% of transcriptome variation, as input to the model.

### Secretory pathway signatures

Boundaries of the secretory pathway were defined using Feizi’s 2017 reconstruction of the mammalian secretory pathway, which consists of 575 core secretory machinery genes divided into 13 subsystems^65^. To extract these secretory machinery genes from our CHO panel, the previously mentioned conversion table ^62^ was used to map CHO genes to their human orthologs. Pathway activity scores were calculated for each of the 13 subsystems using 2 simple equations:

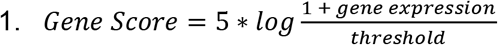

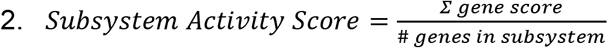

Equation 1 was adapted from thresholding methods implemented in genome-scale model analyses ^28,66^, and involves the preprocessing of the gene expression data using gene-specific thresholds. The threshold is defined by the gene’s mean expression across all samples in the dataset. The activity score is essentially the mean gene score for the subsystem. Activity scores for the 95 recombinant protein expressing CHO cells were normalized to express the change in pathway activity with respect to WT. The relationship between subsystem activity and protein yield were evaluated using non-parametric Spearman correlation. Hierarchical clustering and visualization of the activity scores was achieved using the pheatmap package ^67^ in R. The samples were clustered based on euclidean distance and complete linkage clustering. The relationship between stress response genes and protein yield within cluster 3 were evaluated using non-parametric Spearman correlation. Significance values were adjusted using FDR to correct for multiple testing. Significant correlations were visualized in a lollipop plot using ggplot2 ^57^ in R. The stats package included in R was used to fit a linear model and describe the variance in protein yield explained by genes Derl2 and Alg12.

### Metabolic host response

Expression data from the panel of 96 CHO cultures were subjected to metabolic analysis using CellFie ^28^. CellFie was run using the MT_iCHOv1_final model with the following parameters: local minmaxmean threshold with upper and lower percentile values of 25 and 75 respectively. CellFie provides metabolic task activity in two forms: binary (active or inactive) and quantitative. The binary form of metabolic tasks was used to determine the percent of active vs inactive tasks among the panel of CHO cells. The percent of differentially active tasks was visualized in a boxplot using ggplot2 ^57^ in R. The quantitative form of metabolic task activity was used to calculate differential metabolic activity and correlations with protein yield. Significant differences in metabolic activity between the non producing cell lines and the successfully producing cell lines was performed using a Welch’s t-test on the log2 transformed quantitative activity scores. The relationship between task activity and total protein yield was evaluated using non-parametric Spearman correlation. Task activity that involved specific amino acids were uniquely normalized with respect to the amino acid composition of the protein being expressed in the given sample. Significance values were adjusted using FDRto correct for multiple testing. Significant correlations were visualized in a treemap using the ggtree package^68^ in R.

## Supporting information

Description of Additional Supplementary Files

Supplementary Dat 3

## Acknowledgments

This work was supported by generous funding from NIGMS (R35 GM119850), NIAID (UH2AI153029), the Novo Nordisk Foundation (NNF10CC1016517 and NNF20SA0066621), SSF (SB16-0017), the Swedish innovation agency Vinnova (2021-02640, 2017-02105 and 2016-0518), AstraZeneca and Knut and Alice Wallenberg foundation (Wallenberg Center for Protein Research).

## Author contributions

NEL, JR, MU designed the study and oversaw its implementation and analysis. HOM, CCK conducted the analyses and interpreted the results. MM, ML, AS, HT, SH, AB, LG, DH performed the experiments and collected data. HOM, NEL wrote the manuscript. MM, LG, NEL, JR, HT, CCK, DH critically revised the article.

