## Supplementary Information for "Deciphering the determinants of recombinant protein yield across the human secretome"

**This file contains:**

**Supplementary Results**

**Supplementary Methods**

**Supplementary Figure 1-3**

**Supplementary Table 1**

**Supplementary References**

**Other supplementary materials for this manuscript include the following:**

Supplementary Data 1-13

### SUPPLEMENTARY RESULTS

#### Recombinant protein-specific burdens invoke unique stress response signatures

It is well known that heterologous protein expression can trigger cellular stress response (CSR), a global feedback regulator of protein expression [1]. To investigate the differences in stress response within our panel of CHO cells, we performed differential expression analysis among the subset of genes annotated with the Gene Ontology (GO) term 'cellular response to stress' (GO:0033554) (Fig S3A; Supplementary Data 4; Supplementary Data 11). Of the 1202 stress genes identified in our panel of cells, we identified 118 differentially expressed genes (DEGs) (|FC| >= 2; FDR<=0.01) between the cells that failed to produce protein and the cells that successfully produced protein. We observed a high prevalence of stress response to DNA damage among the DEGs. There is no clear up or down regulation to this response in either group, rather we observe unique responses to this burden. The unique sets of genes activated amongst the two groups alludes to potentially divergent origins or means of coping with DNA damage-induced stressors.

We also observed unique signatures in response to oxidative stress. Oxidative stress occurs when there is an imbalance between antioxidant defenses and the accumulation of oxygen reactive species (ROS), which are known to be generated during recombinant protein production in CHO [2]. Firstly, we noticed that the successfully producing cells show a more profound response to oxidative stress, upregulating almost twice as many oxidative stress response genes compared to the non-producing cells. The ability to mount an adequate response against oxidative stress may facilitate recombinant protein production. Second, we observed that three of the genes depleted in the failed producers encode proteins belonging to the solute carrier (SLC) superfamily, supporting the negative enrichment in SLC transmembrane transport observed in the preliminary GSEA analysis. As a major family of transmembrane proteins responsible for the transport of essential nutrients and metabolites, SLC proteins are critical in many essential physiological functions including oxidative stress. SLC7A11 (solute carrier family 7 member 11) shows the greatest depletion among oxidative stress genes in the failed cells (LFC=-1.85, FDR=5.19E-07) and is involved in the specific transport of cysteine and glutamate. SLC25A24 (solute carrier family 25 member 24), another important SLC transporter which shows depletion in the failed cells, mediates adenosine triphosphate (ATP)-mediated calcium buffering at the mitochondrial matrix, which is essential for normal energy production, protein production within cells, and protecting cells against oxidative stress-induced cell death [3]. Depletion of these SLC transporters could be hindering recombinant protein production. Lastly, several of the oxidative stress response genes upregulated in the producing cells possess functions directly tied to the endomembrane system. For example, GBA encodes a lysosomal membrane protein that functions in the metabolism of complex glycolipids and the turnover of cellular membranes. This could impact overall protein secretion as the secretory pathway is a central component of the endomembrane system.

ER stress response was another prevalent pathway observed among the DEGs. ER stress occurs when there is an imbalance between ER protein folding capacity and protein folding demand leading to an accumulation of misfolded proteins within the ER. In response to ER stress cells activate the unfolded protein response (UPR) which suppresses protein synthesis, induces ER-associated degradation (ERAD) of terminally misfolded protein, and enhances production of protein folding chaperones [4]. Of the few ER stress response genes that showed significant upregulation in the non-producers, half function in ubiquitin-mediated proteasomal degradation suggesting greater protein clearance in the failed producers. Among the many genes significantly depleted in the failed producers were genes involved in all three branches of the UPR: IRE1α (inositol-requiring enzyme-1α), PERK (protein kinase R-like ER kinase), and ATF6α (activating transcription factor 6α). Both PERK (LFC=1.08, FDR=1.15E-13) and ATF6 (LFC=-1.69, FDR=2.14E-36) showed significant upregulation in the successfully producing cell lines. While IRE1 did not meet our strict threshold for significance, it still showed increased expression in the producing cells (LFC=0.89, FDR=5.17E-06) further supported by the significant upregulation (LFC=-2.35, FDR=6.89E-39) of its downstream target EDEM1 (ER degradation enhancing alpha-mannosidase like protein 1). Furthermore, several genes involved in the JNK (c-Jun N-terminal kinase) cascade, another downstream target of the IRE1α UPR branch, were found upregulated in the successfully producing cells compared to the failed producers. Results also showed genes of the heat shock protein family (DNAJC18 and HYOU1) and genes involved in disulfide bond formation (ERO1A and PDIA3) upregulated in the successfully producing cells suggesting greater protein folding capability. Altogether these results suggest that greater activation of ER stress response is associated with more efficient recombinant protein production.

In addition to observing distinct responses to stress between successfully producing cells and the cells that failed to produce any protein, we observed significant correlations between several stress response genes and protein yield (Supplementary Data 5). We identified 13 genes significantly correlated ($|r|>=0.6$; FDR<= 0.05) with protein yield among the subset of cells producing the lowest and highest amount of recombinant protein (Fig S3B). Most stress response genes show a positive correlation with protein yield, further suggesting activation of a proper stress response aids recombinant protein production. Only two genes showed negative correlation with protein yield: SEM1 and RPS27L. SEM1 encodes a subunit of the 26S proteasome involved in proteasome assembly and ubiquitin-binding, again suggesting higher protein turnover in the poorly producing cells. RPS27L is an evolutionarily conserved ribosomal protein that functions in the maintenance of genome integrity and p53-mediated apoptosis [5–8]. Specifically, RPS27L facilitates the repair of DNA interstrand cross-links (ICLs) to maintain genomic integrity [8]. The negative correlation with this gene could indicate higher occurrences of these extremely deleterious ICL lesions in the poorly producing cells.

### SUPPLEMENTARY METHODS

**Host cellular response to stress**

Differential expression between subsets of samples was performed with DESeq2 [66]. The subset of genes involved in cellular response to stress were extracted and annotated using the GO term GO:0033554 'cellular response to stress' retrieved from EMBL-EBI Quick GO database (Supplementary Data 11). Volcano plot visualization of the stress response DEGs was accomplished using the EnhacedVolcano package [67] in R. Visualization of select DEGs and associated GO terms was carried out using the GOplot package [68] in R. The relationship between stress response genes and protein yield were evaluated using non-parametric Spearman correlation. Significance values were adjusted using false discovery rate (FDR) to correct for multiple testing. Significant correlations were visualized using the STRING protein-protein interaction (PPI) network (NDEx UUID:cfcd4cdb-86da-11e7-a10d-0ac135e8bacf) [69] and the ggnet package [70] in R.

### SUPPLEMENTARY FIGURES & TABLES

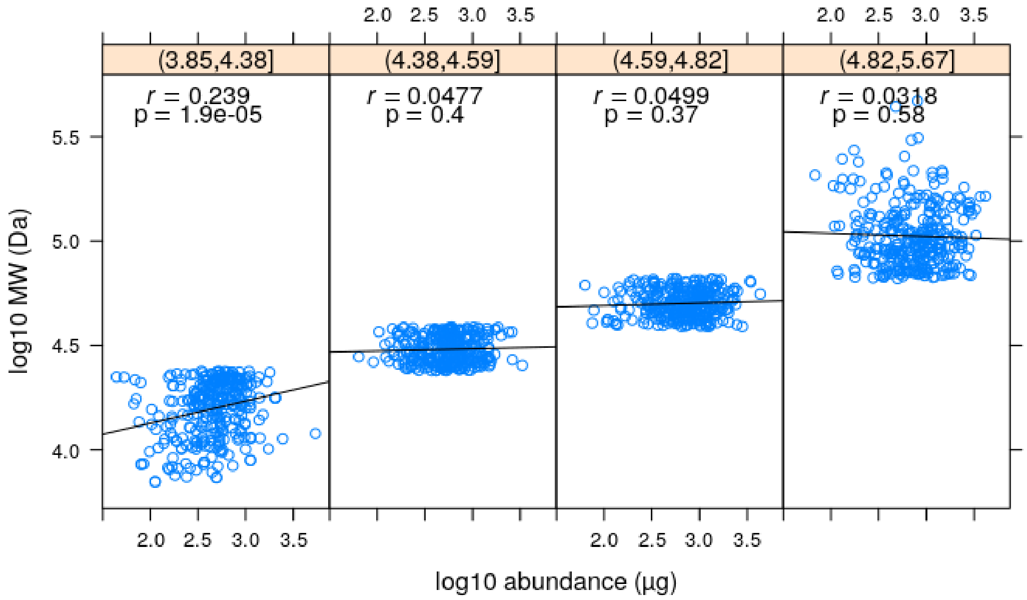


**Fig. S1.** Correlation between protein yield and binned MW. Significant positive correlation only holds true for low MW proteins.

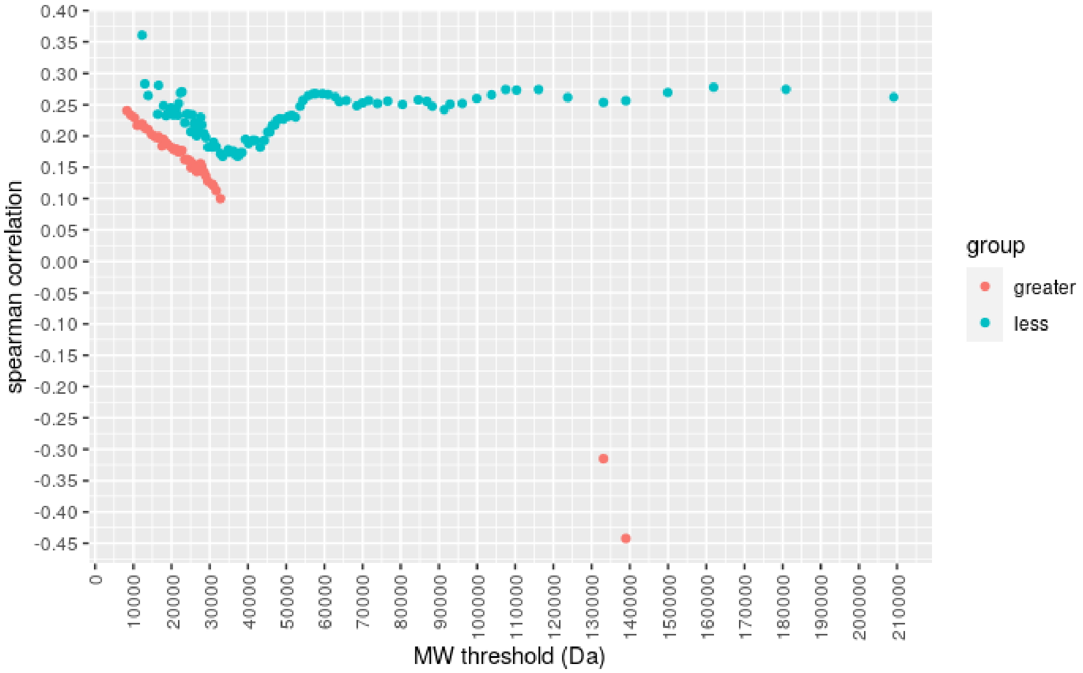


**Fig. S2.** Correlation between protein yield and MW for proteins above and below a series of thresholds. A significant drop in correlation was observed once the protein surpassed 2500-3500 Da.

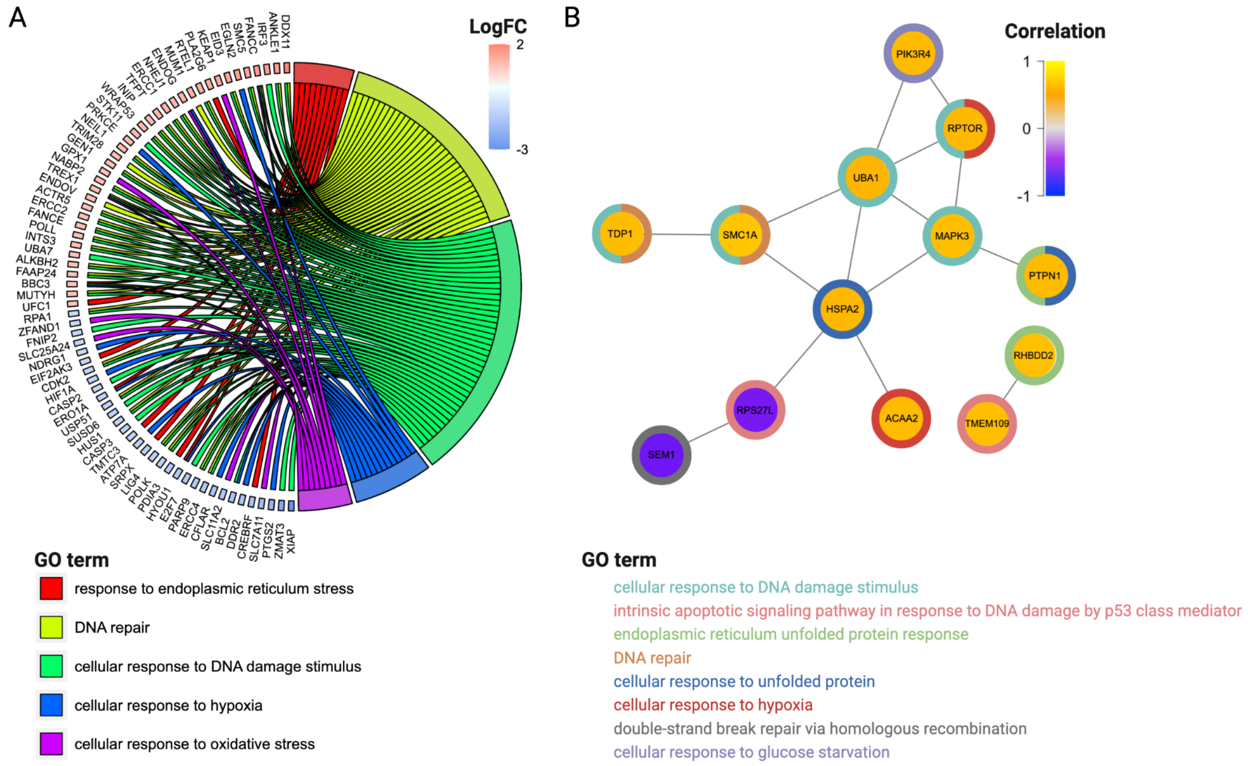


**Fig S3. Stress response cell signatures. A)** Overview showing the relationship amongst GO stress response terms and select significantly differentially expressed genes between the producing and non-producing cells. GO terms have been binned into general categories. **B)** Protein-protein interaction network of the stress response genes which showed significant correlation with protein yield among the high and low producers. Nodes have been colored according to their correlation with protein yield, and node borders have been colored according to select binned GO categories.

**Table S1.** Protein feature's individual correlation with rProtein yield. Cells that failed production were set to 0 titer and included in the analysis.

| **Feature group** *feature* | **Correlation with protein yield** | |
|---|---|---|
| Molecular weight | | |
| *MW (Da)* | 0.055 | |
| AA composition | | |
| *AA.comp R* | -0.190 | *** |
| *AA.comp T* | 0.153 | *** |
| AA composition correlation with CHO | | |
| *AA.comp correlation with essential native CHO* | 0.176 | *** |
| AA class composition | | |
| *AA.comp med volume* | 0.176 | *** |
| *AA.comp basic residues* | -0.170 | *** |
| Post-translational modifications | | |
| *N-linked glycosylation* | 0.260 | *** |
| *Transmembrane domain* | 0.163 | *** |
| Secondary structure | | |
| *Coil* | -0.24 | *** |
| Relative solvent accessibility | | |
| *Mean accessibility score* | -0.13 | * |
| *Percent hydrophobic solvent-inaccessible residues* | 0.175 | *** |
| *Percent hydrophobic solvent-accessible residues* | -0.07 | |
| Stability & solubility | | |
| *Net charge* | -0.19 | *** |
| *Grand average of hydropathicity* | 0.181 | *** |
| *Isoelectric point* | -0.19 | *** |

List of selected protein features amongst predictors with the strongest Spearman correlation coefficient with protein yield. Cell lines that failed to produce protein were set to 0 yield. Signficance values were adjusted using false discovery rate (FDR) method to correct for multiple testing: *P ≤ 0.01, **P ≤ 0.001, ***P ≤ 0.0001.
