## Supplementary material for "Deciphering the determinants of recombinant protein yield across the human secretome": Description of Additional Supplementary Files

**File Name: Supplementary Data 1**
**Description:** RNA-Seq quantification of transgene mRNA abundance for the panel of 95 human proteins expressed in CHO and their total protein yield in micrograms.

**File Name: Supplementary Data 2**
**Description:**

- Description table (sheet 1): Table describing and categorizing the 218 curated protein features used as potential predictors of protein abundance.
- All features (sheet 2): Comprehensive table of protein features for the HSP proteins.

**File Name: Supplementary Data 3**
**Description:** Results of principal component analysis performed on the 96 RNA-Seq cells.

**File Name: Supplementary Data 4**
**Description:** Results from the differential expression analysis among the subset of genes annotated with the Gene Ontology (GO) term ‘cellular response to stress’ (GO:0033554).

**File Name: Supplementary Data 5**
**Description:** Spearman correlation between stress response genes and protein yield among the subset of cells producing the lowest and highest amount of recombinant protein.

**File Name: Supplementary Data 6**
**Description:** Secretory pathway subsystem activity scores for the panel of 95 recombinant protein expressing CHO cells and WT.

**File Name: Supplementary Data 7**
**Description:** Spearman correlation between secretory pathway genes and protein abundance for the cells of cluster 3.

**File Name: Supplementary Data 8**
**Description:** Output from the CellFie metabolic analysis of the panel of 95 recombinant protein expressing CHO cells and WT.

**File Name: Supplementary Data 9**
**Description:** Spearman correlation between metabolic task scores and protein yield among the subset of cells producing the lowest and highest amount of recombinant protein.

**File Name: Supplementary Data 10**
**Description:** Cleaned up version of the Human secretome productivity data taken from Tegel et al 2020 and used in this study.

**File Name: Supplementary Data 11**
**Description:** Genes involved in cellular response to stress extracted from EMBL-EBI Quick GO database using term GO:0033554 ‘cellular response to stress’.

**File Name: Supplementary Data 12**
**Description:** Spearman correlation between secretory pathway genes and protein abundance across the panel of 95 recombinant protein expressing cells.

- Sheet 1 (subsystems): Spearman correlation between the 13 secretory pathway subsystem activity scores and protein abundance across the panel of 95 recombinant protein expressing cells.
- Sheet 2 (genes): Spearman correlation between secretory pathway genes and protein abundance across the panel of 95 recombinant protein expressing cells.

**File Name: Supplementary Data 13**
**Description:** RNAseq (TPM and counts) of the 96 cells.
